# Neuronal tRNA Modifications in *Aplysia californica* are Repatterned during Behavioral Habituation

**DOI:** 10.1101/2025.11.13.688303

**Authors:** Han Xin Huang, Gabriella M. Floro, Emmanuel Asare, Weichen Huang, Kevin D. Clark

## Abstract

Transfer RNA (tRNA) modifications have been increasingly implicated as post-transcriptional regulators of basic neuronal functions. However, characterizing tRNA modification profiles in specific neural circuits that control well-defined behaviors is notoriously difficult due to the complexity of conventional neurobiological models. Here, we leveraged the numerically simple central nervous system (CNS) of the marine mollusk *Aplysia californica* to investigate tRNA modification dynamics in functionally identified neurons during habituation of a defensive reflex. We identified and quantified dozens of neuronal tRNA modifications using liquid chromatography-tandem mass spectrometry (LC-MS/MS), revealing characteristic distributions of select small RNA modifications across different neuronal and non-neuronal tissues. Upon behavioral habituation of the siphon-elicited siphon withdrawal reflex (SSWR), we found that tRNA modification profiles in the major ganglion that controls the SSWR were repatterned with predictable, learning-related changes. We discovered a family of anticodon loop modifications including *N*6-isopentenyladenosine (i^6^A) and its downstream product, 2-methylthio-*N*6-isopentenyladenosine (ms^2^i^6^A), that displayed a significant increase and trend toward lower levels, respectively. These tRNA modification dynamics occurred independently of changes in expression of their parent tRNAs, illustrating that behavioral habituation alters the activity of tRNA-modifying enzymes. Overall, our work reveals an underexplored link between tRNA modifications and behavioral habituation and provides new insights toward understanding the post-transcriptional mechanisms of learning and memory.

**One-Sentence Summary:** Characterization of neuronal tRNA modifications during behavioral habituation in *Aplysia californica* revealed a post-transcriptional mechanism of learning and memory.

**Graphical Abstract:** 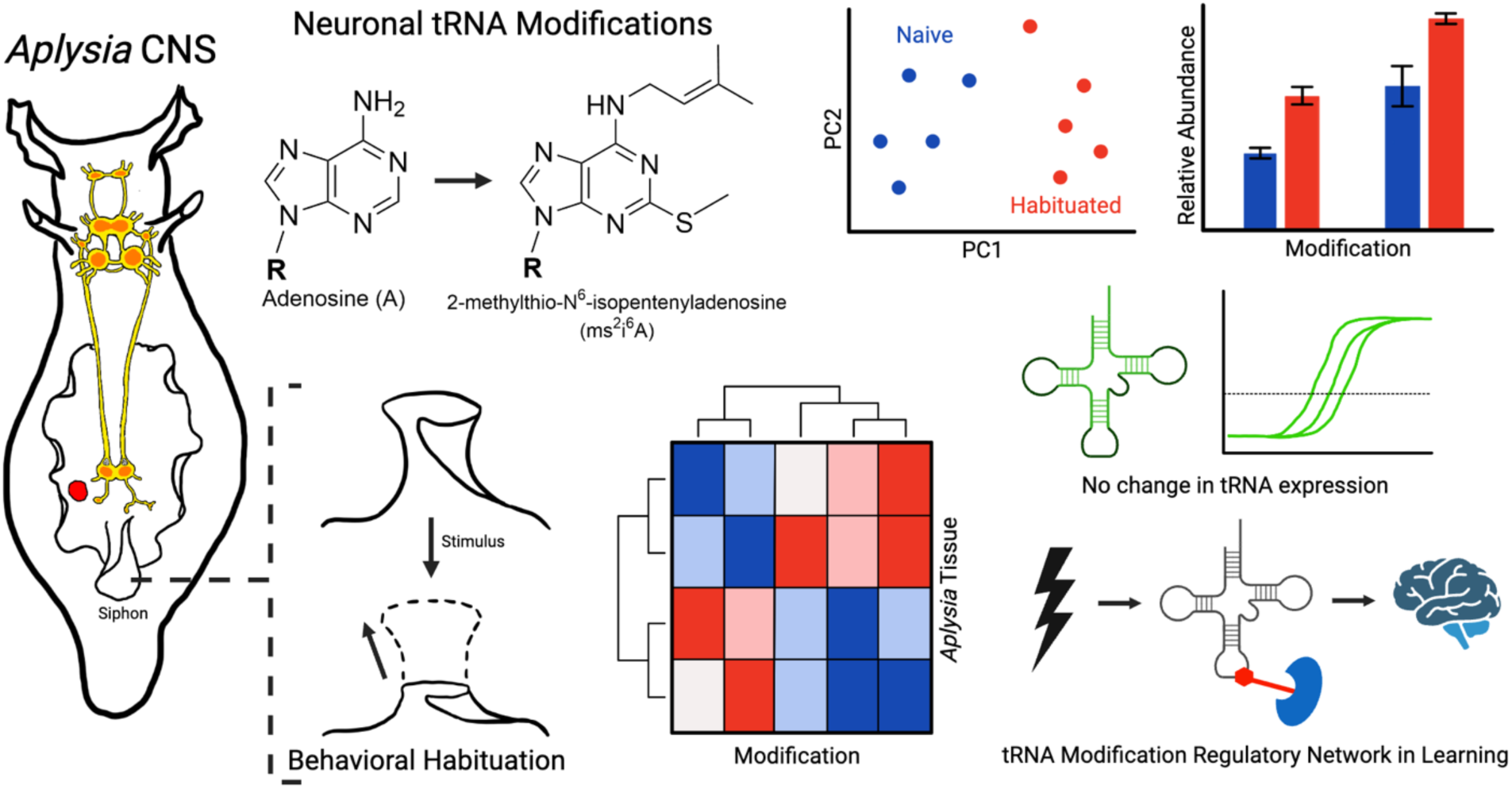

## INTRODUCTION

RNA modifications are enzymatic alterations to canonical nucleotides of RNA that regulate protein translation and a broad array of cellular functions (*1*). To date, 170 post- and co-transcriptional modifications have been discovered that range in complexity from methylations to transglycosylations. Some of these modifications are enzymatically reversible, creating a dynamic epitranscriptome landscape that has been implicated in cell stress response (*2*) and cell-to-cell signaling (*3*). Transfer RNAs (tRNAs) are the most abundantly modified RNA subtype, and their modifications are integral components of the cellular translation apparatus that regulate protein synthesis via accurate codon recognition and reading frame maintenance (*4*, *5*). tRNA abundances and modification stoichiometries are dynamically adjusted in response to environmental cues (*6–10*), which in turn leads to codon-biased translation of proteins that potentiate an appropriate cellular response (*7–10*). In the central nervous system (CNS), stimulus-responsive tRNA modifications may play similar roles and ultimately connect to observable changes in animal behavior. However, the numerical complexity of conventional neurobiological models (10^5^–10^11^ CNS neurons (*11–13*)) precludes characterization of tRNA modification dynamics in well-defined neural circuits that control specific behaviors. While tRNA modifications and their corresponding enzymatic writers and erasers have been linked to neurological diseases and disorders (*14*, *15*), synapse formation (*16*), learning and memory (*5*, *17*), and cognitive impairment (*18*, *19*), correlating tRNA modification dynamics with well-defined CNS functions remains a significant challenge.

*Aplysia californica* is a marine mollusk and classical neurobiological model that is especially useful for studying the molecular underpinnings of fundamental CNS functions (*20*, *21*). The *Aplysia* CNS is comprised of ∼10^4^ neurons, and many of these neurons have been characterized by electrophysiology (*22*) and molecular techniques (*23*) to glean their functional identities (*24*). As a primary example, the specific neural circuits controlling defensive reflexes like the withdrawal of respiratory organs (e.g., the gill and siphon) have been thoroughly characterized in *Aplysia* (*21*). These reflexes are readily modulated by elementary forms of learning including sensitization and habituation, during which repeated exposure to aversive stimuli prolongs or attenuates responses to subsequent innocuous stimuli, respectively (*20*, *21*). These forms of non-associative learning involve release of the monoamine neurotransmitter serotonin (5-HT) and are divided into short- and long-term stages that have distinct molecular hallmarks, including post-translational modification of ion channels and the activity-dependent synthesis of new proteins (*21*, *25–29*). Intriguingly, a specific tRNA anticodon modification, 5-methoxycarbonylmethyl-2-thiouridine (mcm^5^s^2^U), is known to alter the translation of learning related proteins and increase neuronal excitability during behavioral sensitization (*30*). However, little is known about how the full complement of neuronal tRNA modifications impacts animal behavior and whether post-transcriptional mechanisms are generalizable to other forms of learning. In contrast to sensitization, behavioral habituation is characterized by gradually diminishing responses to an applied stimulus and is widely considered one of the simplest forms of learning. Habituation results reduced presynaptic activity in sensory neurons that induces homosynaptic depression at the sensorimotor synapse (*31*, *32*). Enduring forms of habituation require RNA synthesis (*33*), the synthesis of new proteins (*34*), and post-translational modifications (*35*) that are distinct from those required for sensitization. However, the mechanisms that govern protein translation during behavioral habituation remain unresolved (*36*).

In this study, we elucidated the dynamic tRNA modification landscape of the *Aplysia* CNS during behavioral habituation. We used a liquid chromatography-tandem mass spectrometry (LC-MS/MS)-based neuro-epitranscriptomics approach (*30*, *37*) to simultaneously identify and quantify dozens of tRNA modifications across major ganglia. Four modifications, namely, queuosine (Q), mannosyl-queuosine (man-Q), galactosyl-queuosine (gal-Q), and *N*1-methylinosine (m^1^I), were previously unreported in the *Aplysia* CNS. Habituation of the defensive reflex known as the siphon-elicited siphon withdrawal reflex (SSWR) produced characteristic shifts in the tRNA modification profiles of neurons innervating organs involved in SSWR. Significantly decreased levels of m^7^G and m^1^A were observed in neuronal small RNAs following habituation of the SSWR, which directly contrasts with the changes observed during behavioral sensitization (*30*), illustrating that different learning paradigms have different yet predictable tRNA modification dynamics. Moreover, habituation resulted in a significant increase in the tRNA-specific modification *N*6-isopentenyladenosine (i^6^A), along with a corresponding decrease in the abundance of a downstream product, 2-methylthio-*N*6-isopentenyladenosine (ms^2^i^6^A). Expression levels for tRNAs harboring these modifications were constant, demonstrating that tRNA modification dynamics occur independently of changes in tRNA expression during habituation. Altogether, these results bridge a persistent knowledge gap in our understanding of behavioral habituation by linking tRNA modification dynamics to learning, memory formation, and animal behavior.

## RESULTS

### RNA Modifications Show Selective Enrichment in the *Aplysia* CNS with Distinct Spatial Distribution across Neuronal Tissue

Neuronal RNA modifications are often characterized using methods that target a single modification in complex animal models, limiting our knowledge of how the complete epitranscriptome landscape impacts basic CNS function. Here, we used LC-MS/MS to globally profile RNA modifications in *Aplysia* total RNA extracts from major CNS ganglia and heart tissue, with the initial goal of assessing spatial distributions of RNA modifications within and between tissues. Enzymatic digestion of total RNA to nucleosides and subsequent LC-MS/MS analysis identified a total of 26 unique modified ribonucleosides (Fig. 1 and Table S1), four of which, including Q, man-Q, gal-Q, and m^1^I, were previously uncharacterized in *Aplysia* CNS tissue (Fig. S1-S3). We then enriched neuronal small RNAs (< 200 nt) from these tissues and determined the variation in RNA modification abundances between animals. Modified nucleoside peak areas displayed average relative standard deviations (RSDs) of 21.90 ± 21.96% (Table S2), with the buccal and pedal ganglia producing the highest and lowest RSD values of 29.59 ± 18.50% and 12.16 ± 7.28%, respectively. The precision of our measurements is superior to prior inter-animal RSDs reported for modified nucleosides in the *Aplysia* CNS (*30*) and is equal to or better than RSDs reported for other biological samples (*e.g*., human urine, plasma, rat liver) (*38*), giving us confidence in the quantitative capabilities of our method.

**Fig. 1.**
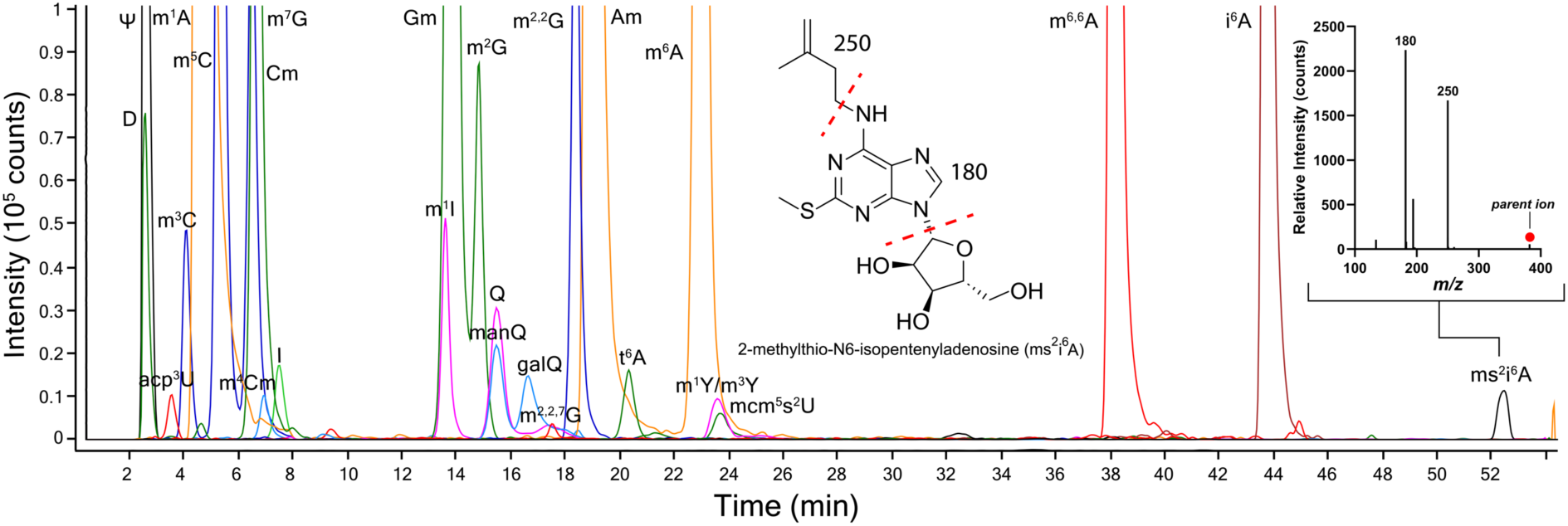
LC-MS/MS characterization of modified ribonucleosides in *Aplysia californica*. Representative overlayed extracted ion chromatograms (EIC) of modified ribonucleosides found in major CNS ganglia and heart. Nucleosides were analyzed following digestion of 4-5 µg of total RNA isolated from neuronal and non-neuronal tissues from a single animal. Panel inset shows MS/MS spectrum of 2-methylthio-*N*6-isopentenyladenosine (ms^2^i^6^A), in which CID fragmentation patterns (shown in chemical structure) enable nucleoside identification.

We then compared tRNA modification abundances across functionally distinct CNS subregions (i.e., major ganglia) and how these profiles compared to a non-neuronal tissue, as it is known that tRNA modification abundances are heterogeneously distributed between tissues in other animals (*39–42*). Fig. 2A shows a depiction of *Aplysia* and its major CNS ganglia and heart that were subjected to small RNA extraction and nucleoside analysis by LC-MS/MS. Hierarchical clustering analysis (HCA) revealed obvious distinctions between small RNA modification profiles in the different tissues (Fig. 2B). Examination of the heatmap showed that the highest relative abundances of modified nucleosides were found in heart tissue compared to the CNS. Among the CNS tissues, the pedal ganglion displayed generally higher relative abundances of RNA modifications. Select nucleoside methylations—m^1^A, Am, m^7^G, and m^3^C—were primary contributors to the observed differences in modification profiles between ganglia. m^1^A and Am were selectively enriched in the buccal and pedal ganglia, respectively, relative to other CNS tissue and the heart (Fig. 2B). m^7^G and m^3^C were similarly enriched in the buccal ganglia compared to other neuronal tissues. These results show that select small RNA modifications are heterogeneously distributed across the *Aplysia* CNS.

**Fig. 2.**
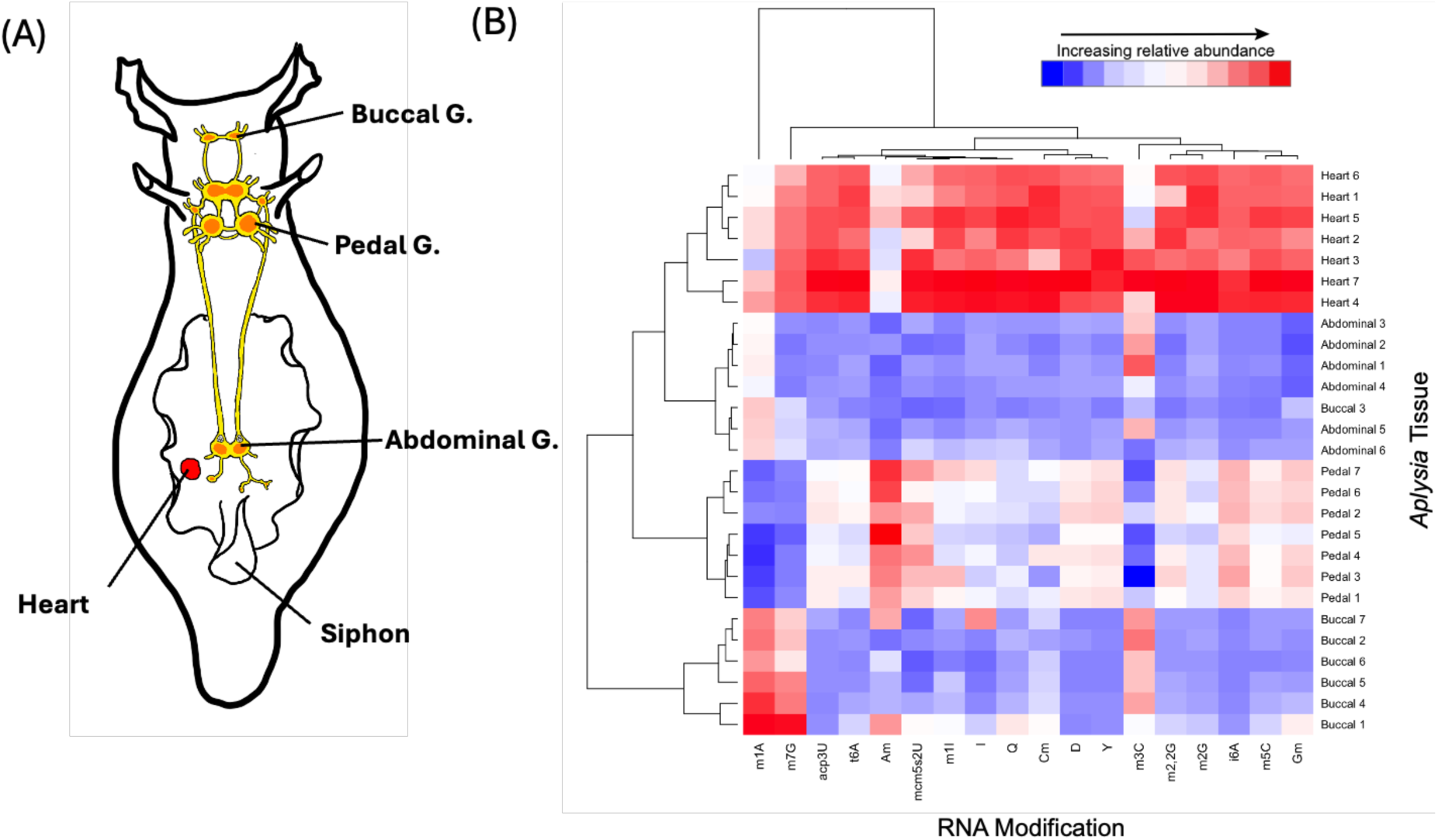
Select small RNA modifications are enriched in the *Aplysia* CNS and show heterogenous distribution across neuronal tissue. (A) Schematic showing major CNS ganglia and organs in *Aplysia*. (B) Heat map diagram of normalized peak areas of modified nucleosides from LC-MS/MS showing hierarchical clustering of tRNA modifications across the abdominal ganglia, buccal ganglia, pedal ganglia, and heart of naïve *Aplysia* (n = 7). Peak areas were normalized by the total peak areas of canonical nucleosides cytidine and uridine.

### Behavioral Habituation in *Aplysia* is Associated with Changes in tRNA Modification Dynamics

Neuronal tRNA modifications have been shown to alter protein synthesis and neuronal excitability during behavioral sensitization in *Aplysia* (*30*), so we asked whether a similar relationship exists between tRNA modifications and a related but distinct form of non-associative learning: habituation. We subjected animals to a massed behavioral training protocol involving mechanical stimulation of the siphon by water jet to facilitate short-term habituation of the SSWR (Fig. 3A). The SSWR initially showed a heightened response to stimulation, followed by a gradual diminution that is characteristic of habituation (Fig. 3B). Trained animals displayed significantly shorter SSWR response times after behavioral habituation (Fig. 3C). We then isolated the abdominal ganglion, which contains sensory and motor neurons that innervate the siphon, immediately following the post-training behavioral test and characterized its small RNA modification profile (Figure S5). The giant R2 neuron, which is not involved in the SSWR circuit and would otherwise contribute ∼10% of the total RNA in the abdominal ganglion, was surgically removed to reduce biological noise (*43*). Unsupervised principal component analysis (PCA) showed noticeable clustering for naïve and habituated animals based on normalized RNA modification levels in the abdominal ganglion (Fig. 3D), with the corresponding loadings plot revealing m^1^A, m^3^C, and m^7^G as primary contributors toward the observed clustering (Fig. 3E). HCA recapitulated the results of behavioral training and PCA, showing general clustering based on behavioral status (Fig. 3F). Notably, levels of m^1^A, m^3^C, and m^7^G markedly decreased in the abdominal ganglion of habituated compared to naïve subjects, while levels of 5-methylcytidine (m^5^C) and *N*6-isopentenyladenosine (i^6^A) increased following habituation. The magnitude of change in the duration of the SSWR between pre- and post-tests did not correspond to nucleoside clustering patterns. We observed high variance due to m^1^A levels relative to other modifications and reasoned that this likely contributed disproportionately to the PCA and HCA clustering results, so we tested whether excluding m^1^A from multivariate analysis would provide insights toward lower abundance modifications that displayed subtler dynamics during the studied learning paradigm. Indeed, clustering on the basis of behavioral status became more obvious when excluding m^1^A from PCA and HCA (Fig. S6). Analyzing the data in this manner reinforced that m^7^G and m^3^C contributed to characteristic small RNA modification dynamics in the abdominal ganglion during learning and that neuronal small RNA modifications in the abdominal ganglion differentiate naïve and habituated *Aplysia*.

**Fig. 3.**
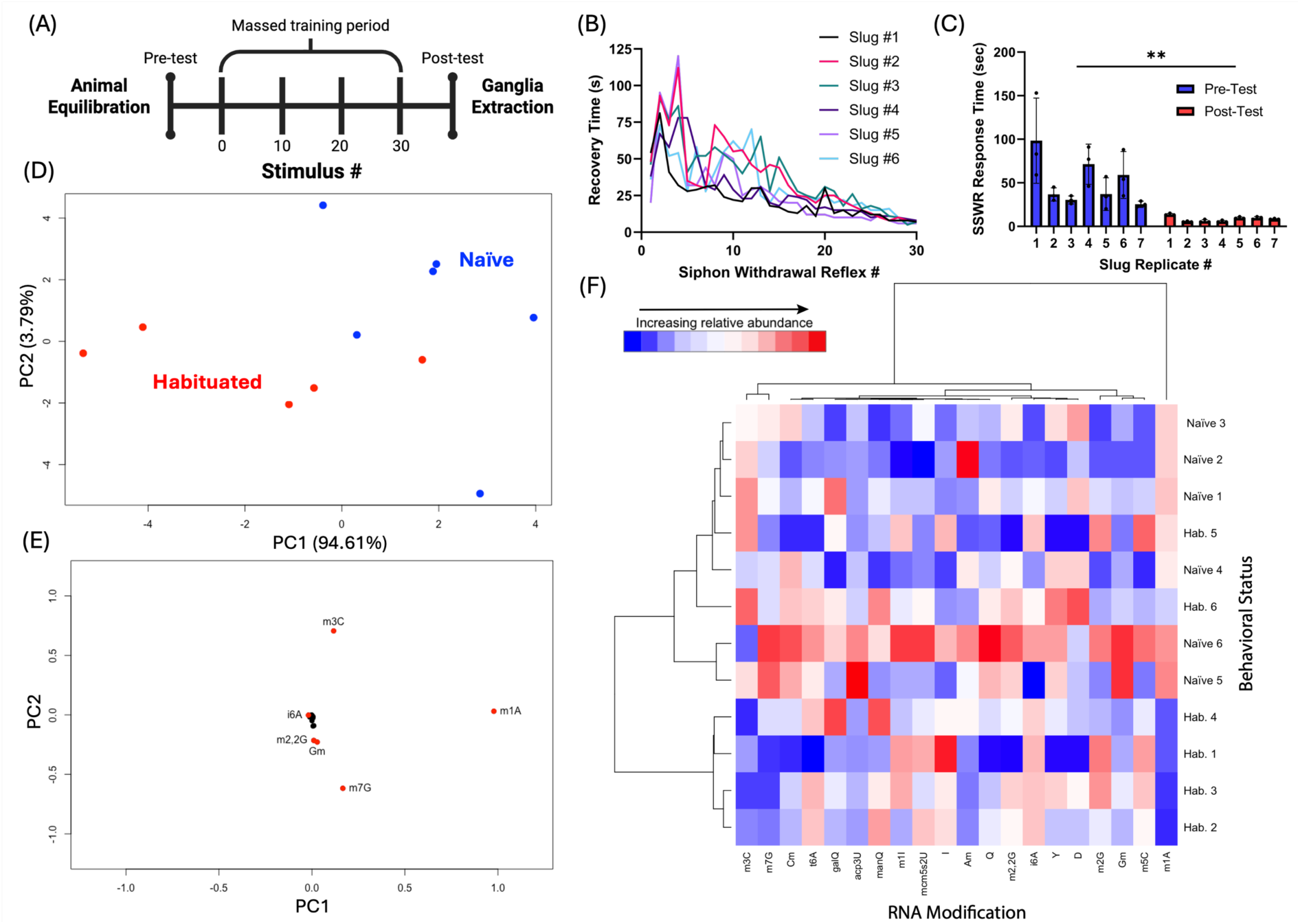
Habituation of the siphon-elicited siphon withdrawal reflex (SSWR) in *Aplysia* is accompanied by changes in tRNA modification profiles within the abdominal ganglion. (A) Behavioral training protocol for habituation of the SSWR in *Aplysia*. (B) Behavioral data for SSWR time plotted against stimulus number (n = 30, inter-stimulus interval = 30 sec). (C) Comparison of SSWR response time of *Aplysia* before and after behavioral training (n = 7). Pre- and post-test measurements were taken in triplicate. (D) Principal component analysis (PCA) of 20 neuronal tRNA modifications detected in abdominal ganglia of naïve and habituated *Aplysia* (n=6) with LC-MS/MS. (E) Corresponding loadings plot with nucleosides selectively labeled based on their contribution towards principal components. (F) Heat map diagram depicting hierarchical clustering in abdominal ganglia of naïve and habituated animals. Paired Student’s t-test: **p<0.005

We then profiled ganglia extrinsic to the SSWR circuit to assess the extent to which the observed small RNA modification dynamics were localized to the abdominal ganglion. We observed subtle clustering for naïve and habituated animals when profiling RNA modifications derived from buccal ganglia, which displayed generally diminished levels of modified nucleosides following training (Fig. S7). We also assessed the small RNA modification profile of the pedal ganglion, which primarily innervates tissues responsible for locomotion (*44*), to determine whether habituation of the SSWR induced global suppression of modification profiles of extrinsic neurons. Both PCA and HCA showed no obvious clustering or trends in small RNA modification patterns between naïve and habituated animals (Fig. S8). Thus, neuronal tRNA modification dynamics associated with habituation of the SSWR are largely localized to the abdominal ganglion.

### Neuronal tRNA Modifications m^7^G, m^3^C, and i^6^A Correlate with Behavioral Habituation

Our untargeted mass spectrometry-based approach revealed select modifications putatively correlated with behavioral habituation. To validate the findings from our initial multivariate analysis, we subjected a new cohort of animals to habituation and quantified a subset of small RNA modifications in the abdominal ganglion that displayed the most obvious changes in our previous experiments, namely, m^1^A, m^7^G, m^3^C, and i^6^A. Pairwise comparisons recapitulated the results from the first cohort of animals, showing significantly decreased levels of m^1^A and m^7^G and increased levels of i^6^A in the abdominal ganglia of habituated animals compared to naïve (Fig. 4A-D). A general but non-significant trend toward lower m^3^C levels was also observed following habituation. Furthermore, we did not observe learning-related dynamics of these modifications in the pedal ganglion or heart tissue.

**Fig. 4.**
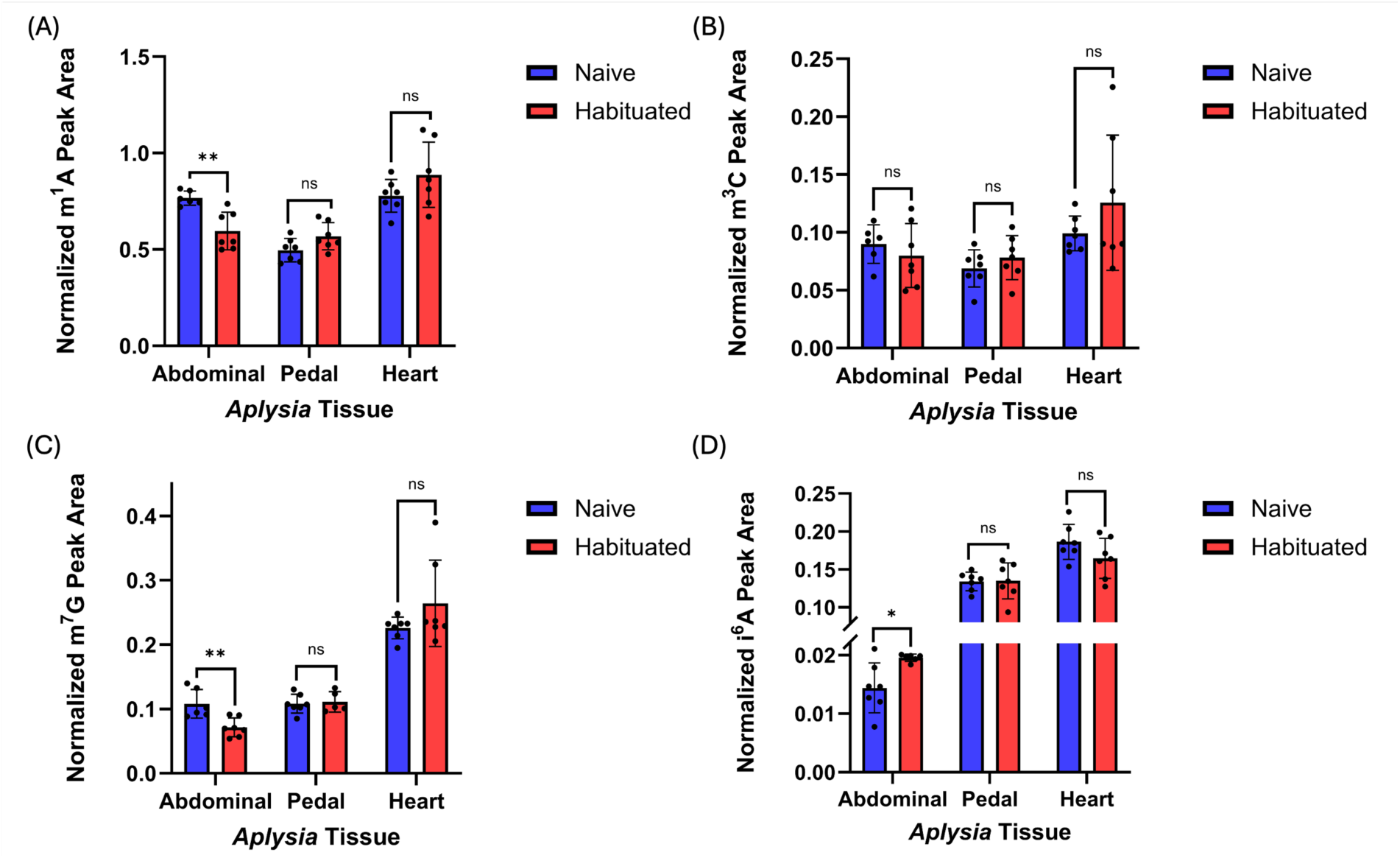
Select small RNA modifications in the abdominal ganglion significantly change following behavioral habituation (n = 7). Pairwise comparisons of normalized (A) m^1^A, (B) m^3^C, (C) m^7^G, and (D) i^6^A levels from LC-MS/MS analysis of small RNAs in different tissues from naïve and habituated animals. Individual peak areas were normalized to the combined peak areas of C and U. Error bars are ± 1SD. Unpaired Student’s t-test: * *p*<0.05 and \*\**p*<0.005.

### Habituation Reprograms i^6^A-family Modifications in Neuronal Mitochondrial tRNAs Independently of tRNA Expression

The significant increase in i^6^A levels we observed in the abdominal ganglion of habituated *Aplysia* prompted us to ask whether the levels of other i^6^A-family modifications were also changing. In mitochondrial tRNAs (mt-tRNAs), i^6^A can undergo a series of subsequent post-transcriptional modifications to form 2-methylthio-*N*6*-*isopentenyladenosine (ms^2^i^6^A) and then 2-methylthiomethylenethio-*N*6-isopentenyladenosine (msms^2^iA) (*45*). ms^2^i^6^A in small RNA fractions from abdominal ganglia showed a strong trend toward lower levels following habituation (Fig. 5A). Notably, 3 out of 10 samples from habituated animals exhibited ms^2^i^6^A levels below the limit of detection of our LC-MS instrument, compared to only 1 out of 10 from naïve animals. msms^2^i^6^A was not detected in tRNA fractions of abdominal ganglia, likely due to its low abundance in the small volume samples.

**Fig. 5.**
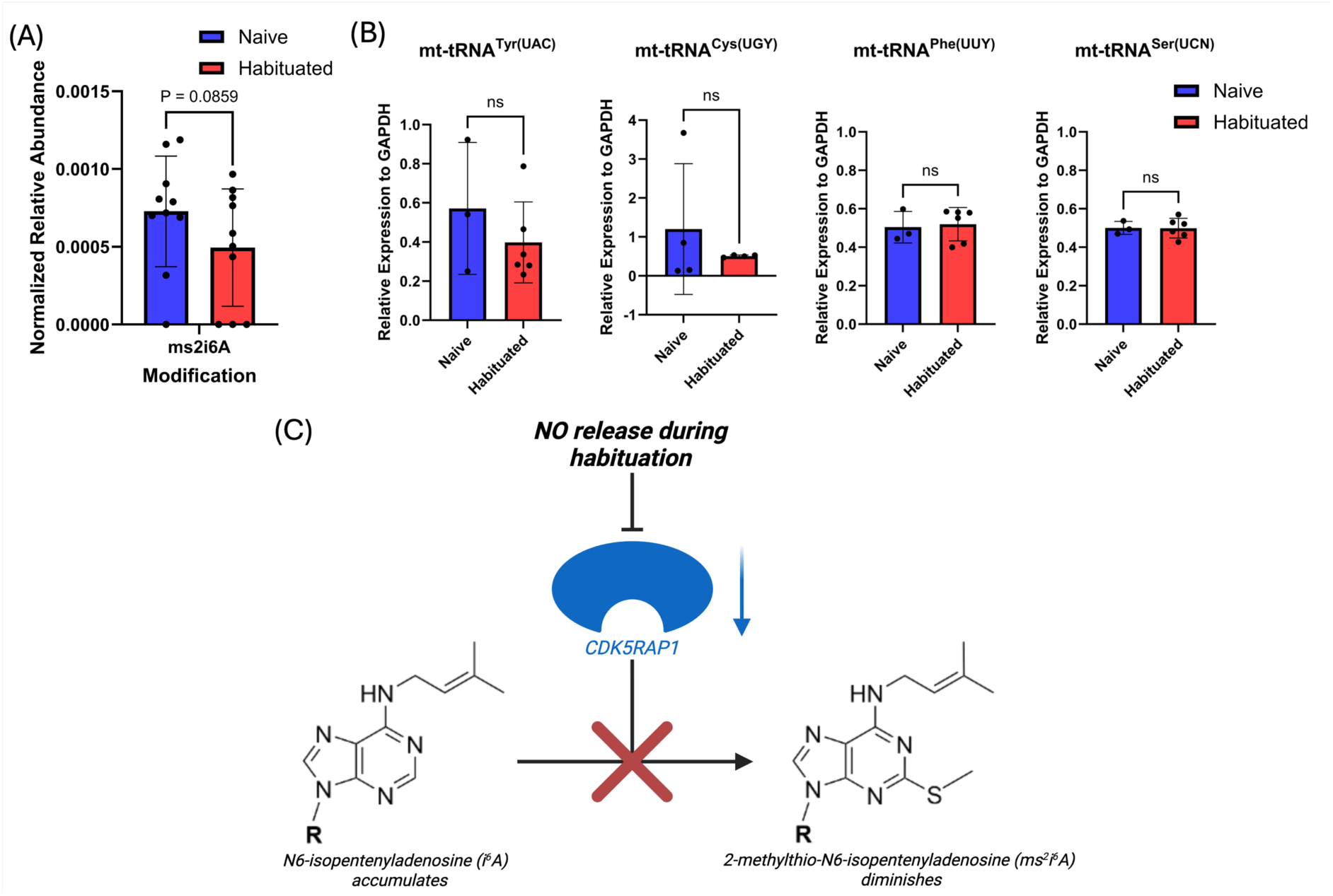
tRNA modification profiles that accompany habituation of the SSWR occur independently of changes in tRNA expression. (A) Comparison of ms^2^i^6^A levels in small RNA fractions from the abdominal ganglion of naïve and habituated *Aplysia* (n = 10, p=0.0859). Relative abundance of ms^2^i^6^A was normalized to the average ΔCq of tRNA transcripts harboring ms^2^i^6^A (-Phe^UUY^ & -Ser^UCN^). (B) Expression levels of tRNA-Tyr^UAC^, - Cys^UGY^, -Phe^UUY^ and -Ser^UCN^ in naïve and habituated animals obtained using RT-qPCR. (C) Proposed mechanistic pathway leading to i^6^A accumulation and ms^2^i^6^A depletion (*49*). Nitric oxide (NO) release during habituation impairs catalytic activity of CDK5RAP1, hindering methylthiolation of i^6^A into ms^2^i^6^A.

We then asked whether the learning-related i^6^A/ms^2^i^6^A dynamics were caused by changes in the expression of their corresponding parent mt-tRNAs. Both i^6^A and ms^2^i^6^A are highly conserved and have been identified at position A_37_ in the anticodon loop of mt-tRNA-Ser^UCN^ and -Tyr^UAC^ (*46*). ms^2^i^6^A has additionally been identified in mt-tRNA-Cys^UGY^ and -Phe^UUY^. While these mt-tRNA modifications have not been mapped in *Aplysia*, they are highly conserved in yeast, humans, and a wide variety of vertebrates and invertebrates (*47*). We thus conducted RT-qPCR experiments using primers specific to tRNA-Tyr^UAC^, -Cys^UGY^, -Phe^UUY^ and -Ser^UCN^ to assess changes in the expression levels of these tRNA transcripts during habituation. No significant changes in the normalized expression levels of these mt-tRNAs were observed in the abdominal ganglia of naïve and habituated animals (Fig. 5B). These results suggest that changes in tRNA modification profiles associated with habituation occur due to changes in the activity of tRNA-modifying enzymes, rather than the expression of new tRNAs. Notably, only transcripts of tRNA-Phe^UUY^ and -Ser^UCN^ showed expression levels with high precision in our experiments; those for tRNA-Tyr^UAC^ and -Cys^UGY^ displayed large variation. This likely results from codon usage bias in the *Aplysia* mitochondrial genome, where certain mRNA codons are favored during translation such that their cognate tRNAs are preferentially expressed (Table S4) (*48*).

## DISCUSSION

Neuronal tRNA modifications in *Aplysia* display characteristic profiles that affect the synthesis of learning-related proteins during behavioral sensitization (*30*), raising the question of whether the roles of RNA modifications are generalizable to other forms learning. Habituation is regarded as one of the simplest forms of learning, yet the promise of unraveling its molecular mechanisms remains unfulfilled. To advance toward this goal, we leveraged the marine mollusk and classical neurobiological model, *Aplysia californica*, and an untargeted LC-MS/MS method to investigate the post-transcriptional mechanisms of behavioral habituation. Our strategy was to first globally profile modified ribonucleosides from small RNA extracts of major ganglia across the *Aplysia* CNS and non-neuronal tissue to assess the distribution of RNA modifications in this animal. These initial experiments revealed that RNA modifications were enriched in select ganglia (Fig. 2B), providing evidence for distinct epitranscriptomic profiles in functionally different cell clusters. Since tissue-specific tRNA modifications are known to display patterns that coincide with different translational requirements (*41*, *49*), our results suggest that distinct tRNA modification profiles within tissue subregions are tuned to meet functional demands of each ganglion.

We then characterized small RNA modification dynamics during behavioral habituation. Whereas sensitization results in an enhanced behavioral response to stimulation, habituation to a stimulus is manifest as a progressively diminishing response. Our results showed that the abdominal ganglia of habituated animals displayed small RNA modification profiles distinct from naïve animals (Fig. 3A-F). The directions of the observed changes in modification levels for naïve and trained animals were largely opposite to those previously observed in sensitized animals (*30*). For example, levels of methylated purine nucleosides m^1^A and m^7^G decreased significantly in habituated animals, compared to significant increases previously observed during behavioral sensitization (*30*). Pairwise comparisons of m^1^A and m^7^G in a separate cohort of naïve and habituated animals validated our findings from multivariate analysis, displaying significantly decreased levels (Fig. 4A, C). Hypomethylation of m^1^A and m^7^G sites in both mitochondrial and cytosolic tRNAs is known to impair neuronal translation (*50*, *51*), raising the question of whether habituation proceeds through post-transcriptional attenuation of global translation or the translation of select proteins. Interestingly, both m^1^A and m^7^G methylations impart a positive charge to the nucleobase, which changes the chemical topology of the RNA and is linked to RNA partitioning to granules (*52*, *53*). In *Aplysia*, stimulus-specific formation RNA-protein granules is a known feature of behavioral sensitization,(*54*, *55*) but further investigation is needed to elucidate whether tRNA m^1^A and m^7^G levels affect granule formation. We also noticed a global decrease in small RNA modifications in the buccal ganglion, which may be explained by a general suppression of buccal ganglion-controlled feeding responses (*56*, *57*) that accompanies non-associative learning in *Aplysia* (*58*). This suggests that consecutive application of noxious stimuli to the siphon can produce concurrent neurochemical changes in neurons extrinsic to defensive reflex circuits, which could be verified by simultaneously gathering data on *Aplysia* feeding behaviors during habituation.

Our LC-MS analysis of small RNA extracts also revealed that habituation significantly increased levels of the tRNA-specific, anticodon loop modification i^6^A in the abdominal ganglion (Fig. 4D). These changes were not observed in the pedal ganglion or non-neuronal tissue, linking tRNA modification dynamics to a specific behavior controlled by the abdominal ganglion. The i^6^A modification is a precursor to several downstream modifications including ms^2^i^6^A, which we found trending to lower levels in neuronal tRNAs from habituated animals. The relatively low abundance of ms^2^i^6^A resulted in some data points falling below the limit of quantification for our LC-MS method, but future development of more sensitive methods for quantifying tRNA modifications is anticipated to enable improved measurements of single *Aplysia* neurons within the SSWR circuit with reduced biological noise and enhanced specificity—progress of which has already been made for certain larger neurons (*59*, *60*). Nonetheless, our LC-MS results suggested that behavioral habituation was accompanied by changes in tRNA expression, the activity of tRNA-modifying enzymes, or a combination of both mechanisms. tRNA expression results showed that the levels of tRNAs known to harbor i^6^A and/or ms^2^i^6^A (mt-tRNA-Ser^UCN^, -Tyr^UAC^, -Cys^UGY^, and -Phe^UUY^) did not change during habituation. These results strongly suggest that the corresponding changes in tRNA modification profiles we observed during habituation were due to modification of pre-existing tRNAs. Since ms^2^i^6^A has no known erasers, the accumulation of i^6^A and corresponding decrease in ms^2^i^6^A levels indicate that habituation is likely the result of reduced activity of an enzymatic writer. Coupled with the observed decreases in m^1^A and m^7^G levels in small RNAs, our findings point to reduced neuronal translation efficiency—largely localized in the mitochondria—as a potential upstream regulatory mechanism of altered CNS function.

The learning-related i^6^A/ms^2^i^6^A dynamics we observed raised two questions: (i) how is the activity of the ms^2^i^6^A writer enzyme modulated by habituation? and (ii) what are the downstream effects of elevated i^6^A/diminished ms^2^i^6^A? CDK5RAP1 is a radical *S*-adenosylmethionine, iron-sulfur cluster enzyme responsible for catalyzing the methylthiolation of i^6^A_37_ to form ms^2^i^6^A_37_ in eukaryotic mt-tRNAs (*61*, *62*). It is conceivable that the co-variation of i^6^A and ms^2^i^6^A we observed by LC-MS is a consequence of changes in the catalytic activity of CDK5RAP1 during habituation that reduce levels of ms^2^i^6^A (Fig. 5C). Nitric oxide (NO) is a well-known inhibitor of iron-sulfur cluster-containing proteins that acts via competitive displacement of their amino acid side chain ligands used for iron-binding to impair catalytic activity (*63*, *64*). Intriguingly, nitric oxide (NO) is also a well-known neuromodulator that is released during behavioral habituation (*65–67*), and has been shown to be necessary for long-term sensitization in *Aplysia* (*68*). Since CDK5RAP1 uses two [4Fe-4S] clusters as cofactors for sulfur insertion into i^6^A_37_ in tRNAs, one might anticipate this writer enzyme to be susceptible to inhibition by NO (*62*, *69–71*). Indeed, NO has been shown to inhibit the bacterial homolog of CDK5RAP1, MiaB, which resulted in higher i^6^A and lower ms^2^i^6^A levels (*72*). CDK5RAP1 responds to cellular redox, where hypoxic conditions altered its activity, resulting in elevated ms^2^i^6^A levels on mitochondrial tRNAs without any changes in mt-tRNA expression (*73*). These findings reinforce a role of reactive oxygen species like NO as upstream modulators of CDK5RAP1 function. It is thus conceivable that increased NO levels released during habituation impair CDK5RAP1 activity, establishing characteristic profiles of i^6^A and ms^2^i^6^A and potentially regulating the translation of select learning-related proteins (Fig. 5C). Given the mitochondrial localization of ms^2^i^6^A-containing tRNAs (*62*, *74*), our results raise the intriguing possibility that the translation of mitochondrial proteins is selectively modulated by mt-tRNA modifications during learning (*62*). The presence of i^6^A-related modifications in tRNAs that decode UNN codons (*75*), as well as evidence of ms^2^i^6^A deficiency in tRNA-Phe and tRNA-Leu modulating proofreading of poly-U codons (*76*), make it plausible that their dynamics can alter the translation of UNN-containing mRNAs that are enriched during learning via codon-biased translation. This work sets the stage for follow up experiments, such as quantitative proteomics and NO synthase inhibition assays, that would clarify the relationship.

In summary, our findings reveal that neuronal tRNA modifications are reprogrammed in response to stimulation during a learning and memory paradigm, raising new, testable questions about the molecular mechanisms underlying behavioral habituation. For example, follow-up quantitative proteomics experiments would reveal how the dynamics of anticodon loop modifications i^6^A and ms^2^i^6^A observed here affect codon-biased translation of mitochondrial proteins. Finally, leveraging the *Aplysia* model for measuring tRNA modifications in single, functionally identified neurons of the SSWR circuit is sure to reduce biological noise from extrinsic neurons and enable us to distinguish subtler learning-related RNA modification dynamics.

## MATERIALS AND METHODS

### Reagents & Materials

Ammonium bicarbonate, bovine serum albumin, kanamycin sulfate, and magnesium chloride were purchased from Sigma-Aldrich (St. Louis, MO, USA). Acetonitrile (LC-MS grade), ammonium acetate (LC-MS grade), chloroform, ethanol, isopropanol, Trizol reagent, water (LC-MS grade), DNase I RNase-free kit, RiboLock RNase Inhibitor (40 U/µL), and Invitrogen Superscript IV Reverse Transcriptase kit were obtained from Thermo Fisher Scientific (Waltham, MA, USA). Monarch Spin RNA Cleanup Kit (50 μg, #T2040L), nuclease P1, snake venom phosphodiesterase, and bacterial alkaline phosphatase were purchased from New England BioLabs Inc. (Ipswich, MA, USA). 100 nmol, standard desalted, single stranded DNA oligonucleotide primers were ordered from Integrated DNA Technologies (USA). SsoAdvanced Universal SYBR Green Supermix was purchased from BioRad (USA). Reagents were used without further purification.

### Animals and Behavioral Training

*Aplysia californica*, weighing ∼100-250 g, were obtained from the National Resource for *Aplysia* (Miami, Florida, USA). Animals were housed in an aquarium with circulating, aerated artificial seawater (ASW) at 14 °C for at least 24 h prior to experimentation. Behavioral training followed a previous experimental design (*30*) with slight modifications to achieve animal habituation. To prepare for behavioral training, animals were placed in ASW-filled isolation tanks and acclimated for 15-30 min. To habituate the SSWR, animals were subject to a massed training protocol comprising of 30 consecutive stimuli delivered to the animal’s siphon with an interstimulus interval (ISI) of 30 s. Stimulus was applied mechanically through an electric water-flosser (Water Pik, Inc., Fort Collins, CO, USA) at the lowest pressure setting, with the tip placed ∼1 cm away from the siphon upon stimulation. Pre- and post-test evaluations of the SSWR were conducted using the aforementioned procedure before and after behavioral training, respectively. The duration of the SSWR time was measured from the moment the siphon was withdrawn beneath the parapodia upon stimulation to the moment the siphon breached the parapodia upon relaxation, after which the ISI timer began. Individuals that did not exhibit a noticeable withdrawal reflex and recovery were excluded from the study. Prior to dissection, animals were anesthetized by injection with chilled 0.33 M MgCl_2_ (approximately 40% body volume) directly into the body cavity. Relevant CNS ganglia were surgically isolated and stored in ASW before total RNA extraction. Naïve animals were similarly placed in isolation tanks for the duration of the training period to eliminate animal housing differences across treatment groups.

### Ganglion Preparation, RNA Extraction, and Enzymatic Digestion

The abdominal, pedal, and buccal ganglia and heart were surgically isolated from the *Aplysia* CNS from naïve and habituated animals (*59*, *60*). The abdominal ganglion was then immediately subject to protease treatment (10 mg/ml in an ASW solution containing 100 ppm kanamycin antibiotic) for 45 min at 35 °C. The ganglion was then transferred to a Petri dish containing chilled ASW for surgical removal of the R2 neuron to reduce biological noise (*77*). A trimmed transfer pipette, lubricated with a ∼1 mg/mL bovine serum albumin solution, was used for ganglion transfer. Following removal, the ganglion was placed in cold TRIzol reagent (200 µL for abdominal and buccal ganglia; 300 µL for pedal ganglia and heart). All other ganglia and the heart were individually kept in ASW at 4 °C during protease treatment and cell isolation and were placed in TRIzol simultaneously with the abdominal ganglion (*37*).

Following placement into TRIzol, *Aplysia* CNS and heart samples were homogenized using a pestle to ensure cell lysis (*37*). Samples were then mixed with chloroform (30% volume of TRIZol), vortexed, and centrifuged at 15,000*g* at 4 °C for 5 min, where the aqueous supernatant was subsequently isolated. Chloroform extraction was repeated on the aqueous supernatant to ensure thorough removal of organic contaminants, after which the crude RNA extracts were suspended in isopropanol (IPA) overnight at -20 °C.

Total RNA extracts in IPA were centrifuged at 16,200 x *g* at 4 °C for 15 min, where the resultant supernatant was removed to waste. The precipitated RNA pellet was then resuspended in 70% ethanol, centrifuged at identical settings, and the supernatant was removed. After two washes, the RNA pellet was left covered in a fume hood for 2-3 h, or until all supernatant has evaporated. Purified total RNA samples were finally resuspended in nuclease-free water in preparation for small RNA fractionation.

Total RNA samples were subject to small RNA fractionation using the Monarch Spin RNA Cleanup Kit according to the manufacturer’s instructions with slight modifications (*78*). The small RNA eluate was collected, washed, and spun twice to maximize sample purity and yield. Resulting fractions largely consisted of highly structured small RNA (<200 nt), but sufficient residual mRNA to use GAPDH as a housekeeping gene in RT-qPCR experiments. Small RNA samples were eluted in 50 µL nuclease-free water and tested for RNA concentration and purity using a DeNovix spectrophotometer. To align sample volumes for enzymatic digestion, the samples were placed in a vacuum centrifuge for 1 hr at 43 °C, or until all solvent had evaporated. Dried small RNA samples were resuspended in 15 µL nuclease-free water for enzymatic digestion.

Purified small RNA samples were enzymatically digested using a two-step digest protocol (*79*). Samples containing 0.5-2 µg of tRNA were first incubated in 10 mM ammonium acetate (pH 5.0) with 0.6 U Nuclease P1 at 45 °C for 2 hr, and then in 0.1 M ammonium bicarbonate (pH 8.0) with 0.0025 U snake venom phosphodiesterase and 0.01 U bacterial alkaline phosphatase at 37 °C for 2 hr. Samples were then placed in autosampler vials for LC-MS analysis.

### Analysis by LC-MS/MS

Digested small RNA samples were analyzed using an Agilent 1260 Infinity II LC System with diode array detection (Agilent Technologies, Santa Clara, CA, USA) coupled to an Agilent 6530B quadrupole-time-of-flight mass spectrometer using dual Agilent jet stream electrospray ionization operated in positive mode. LC separations were conducted using a ZORBAX Eclipse Plus C18 Narrow Bore RR 2.1 x 100 mm, 3.5 µm column (Agilent Technologies). Gradient elution was performed using a mobile phase A of 5 mM ammonium acetate (pH 5.4) and a mobile phase B of 60/40 mixture of 5 mM ammonium acetate (pH 5.4) / acetonitrile, with a flow rate of 0.2 mL / min at the following conditions: 1% B from 0-5 min, 5% B at 9 min, 7% B at 11 min, 10% B at 13 min, 15% B at 32 min, 70% B at 40 min, 100% B at 44 min with a 10 min hold at 100% B, and a 15 min hold at 1% B prior to the subsequent injection. Mass spectrometry (MS) data collection was conducted using a mass range of 100-3200 *m/z* at a scan rate of 1 spectra/s, The capillary, nozzle, and fragmentor voltages were set at 3000 V, 1000 V, and 100 V, respectively, and the N_2_ drying gas was set to 5 L/min at 300 °C. MS/MS acquisition was performed with a mass range of 100-600 *m/z* at a scan rate of 1 spectra/sec. Precursor ions were selected for fragmentation using an autonomous method based with specific absolute intensity thresholds for collision-induced dissociation of a specific ion and confirmed using the MODOMICS database (*80*).

### LC-MS Data Analysis

The identity of each nucleoside was determined by comparing the dominant MS/MS fragment ions to those reported in the MODOMICS database for compounds eluting at appropriate LC retention times relative to the canonical nucleosides. Individual peak areas were obtained using MassHunter Qualitative Analysis 10.0 & Quantitative Analysis software (Agilent Technologies) from extracted ion chromatograms that correspond to the detected mass of the ionized nucleoside [M+H]^+^ within a qualifier window of 20 parts-per-million (ppm). A broader range of up to 50 ppm was used to accommodate select, low-abundance nucleosides in fractions containing relatively less RNA (< 0.5 ug). For isomers with identical monoisotopic masses and fragmentation patterns (*e.g*., man-Q and gal-Q), identification was based on known elution order from prior separation data using reverse-phase HPLC (RP-HPLC) (*81*). To accommodate variations in the quantity of RNA subject for LC-MS analysis, peak areas of modified nucleosides were normalized relative to the sum of canonical cytidine and uridine peaks while ensuring that canonical signal intensities did not exceed the saturation limit of the MS detector (*82*, *83*). Principal component analysis (PCA) was performed in RStudio (*84*), using a data matrix containing normalized peak area for modified nucleosides with a signal-to-noise ratio > 3. Score and loading plots were generated for the first two principal components. RStudio was also used to conduct unsupervised hierarchical clustering analysis, with Euclidean distance and visualization by heatmap. An unpaired, two-tailed Student’s t-test with equal variance was used to determine significant differences in select RNA modification levels following facilitation of a learning paradigm in the animals.

### tRNA Expression Analysis with RT-qPCR

Extracted small RNA samples from tissue of naïve and habituated *Aplysia* were standardized to 10 ng/µL each with nuclease-free water and subject to DNAse treatment to eliminate genomic DNA contaminants. DNAse treatment was performed using the DNAse I, RNAse-Free kit. DNAse-treated samples were run on the CFX Duet Real-Time PCR System (Bio-Rad Laboratories, Waltham, MA, USA) prior to reverse transcription to assess treatment efficiency, where absence of amplification suggested successful removal of genomic DNA.

DNAse-treated tRNA samples were reverse-transcribed using the Superscript IV Reverse Transcriptase kit according to the manufacturer’s instructions with 2 μM gene specific primers (Table S3). Corresponding cDNA was obtained using reverse primers corresponding to various tRNA isodecoders and the reference gene GAPDH. Primer sequences were designed according to the *Aplysia californica* NCBI RefSeq Assembly (GCF_000002075.1, AplCal3.0) using the *Aplysia californica* mitochondrion, complete genome (NCBI Reference Sequence: NC_005827.1), which was obtained from NCBI Genome Project.(*85*) Primer melting temperatures (Tms) were determined using the OligoAnalyzer Tool from IDT (Integrated DNA Technologies). Primers were also ordered and synthesized from IDT.

RT-qPCR amplification and analysis were performed using the CFX Duet Real-Time PCR System and the CFX Maestro software v2.3 (Bio-Rad), respectively. Cycle quantification (Cq) values were determined according to the ‘Single Threshold Method’ in the software. Samples were prepared for qPCR using the SsoAdvanced Universal SYBR Green Supermix (Bio-Rad) according to the manufacturer’s directions. A two-step reaction was performed with the following thermal cycling protocol: initial denaturation for 3 min at 94 °C, 40 cycles of 10 s at 95 °C, and 10 s at a temperature gradient between 55.8 °C and 57.9 °C. Fluorescence signal was measured at the end of each annealing step. Melting curve analysis was conducted with a temperature gradient of 0.5 °C/s from 60-90 °C to confirm selective amplification of desired products.

Relative expression of tRNAs was performed by normalizing the expression of individual transcripts against that of the housekeeping gene GAPDH to yield ΔCq. A normalized ΔCq for each sample was then obtained via division by the average ΔCq amongst biological replicates for each tRNA transcript (n = 6) (Eq. 1). From a scale of 0 to 1, a higher normalized ΔCq suggested higher relative expression of the target transcript relative to GAPDH, and vice versa. Equations 1 and 2 were used for appropriate calculation of relative expression.

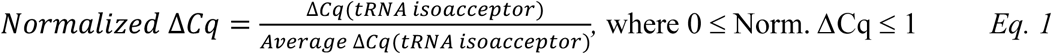

## Supplementary Materials

Fig. S1. Identification of queuosine (Q) in total RNA extracts from neuronal tissue of *Aplysia* using LC-MS/MS.

Fig. S2. Identification of mannosyl-queuosine (man-Q) and galactosyl-queuosine (gal-Q) in total RNA extracts from neuronal and non-neuronal tissue of *Aplysia* using LC-MS/MS.

Fig. S3. Identification of 1-methylinosine (m1I) in total RNA extracts from neuronal and non-neuronal tissue of *Aplysia* using LC-MS/MS.

Fig. S4. Identification of N4,2-O-dimethylcytidine (m4Cm) in total RNA extracts from neuronal and non-neuronal tissue of *Aplysia* using LC-MS/MS.

Fig. S5. Experimental workflow for behavioral training and RNA extraction in *Aplysia.* Fig. S6. Removal of m1A from multivariate analysis improves unsupervised clustering between naïve and habituated *Aplysia*.

Fig. S7. Neuronal tRNA modifications in buccal ganglia subtly distinguish between naïve and habituated *Aplysia*.

Fig. S8. Neuronal tRNA modifications in pedal ganglia do not distinguish between naïve and habituated *Aplysia*.

Table S1. Successfully characterized modified ribonucleosides in 5 major ganglia and heart of *Aplysia californica* (n = 12).

Table S2. Statistical analysis of relative peak areas of modified nucleosides detected by LC-QTOF-MS/MS from small RNA extracts of abdominal (n = 6), pedal (n = 7) ganglia and the heart (n = 7) of *Aplysia*.

Table S3. Target mitochondrial tRNAs and their corresponding forward and reverse primer sequences.

Table S4. Relative codon usage of target tRNAs in *Aplysia* mitochondrial genome. Data file S1. MDAR Checklist

Data file S2. Code Availability for Multivariate Analysis

## Acknowledgements

Funding: KDC acknowledges support from the Arnold and Mabel Beckman Foundation through a Beckman Young Investigator Award.

## Author Contributions

Conceptualization: KDC Methodology: HH, G.F, EA, KDC Investigation: HH, GF, WH Visualization: HH, GF

Funding acquisition: KDC Project administration: HH, KDC Supervision: KDC

Writing – original draft: HH

Writing – review & editing: HH, GF, KDC

**Competing Interests:** The authors declare that they have no competing interests.

**Data and Materials Availability:** MS data for newly detected and established nucleosides are available in the Supplementary Materials. All other data needed to evaluate the conclusions in the paper are present in the paper or the Supplementary Materials.

**Code Availability:** Code used for RStudio data analysis can be found in the document *Code Availability for Multivariate Analysis* in the Supplementary Materials. Instructions for the code are detailed in the document and in the Materials and Methods section *LC-MS Data Analysis*.

