## Supplemental file for "Neuronal tRNA Modifications in *Aplysia californica* are Repatterned during Behavioral Habituation"

**Supplementary Materials for**
**Neuronal tRNA Modifications in *Aplysia californica* are Repatterned during Behavioral Habituation**

Huang *et al*.

**This PDF file includes:**

Figures S1-S8

Tables S1-S3

**Other Supplementary Materials for this manuscript includes the following:**

MDAR Reproducibility Checklist
Code Availability for Multivariate Analysis

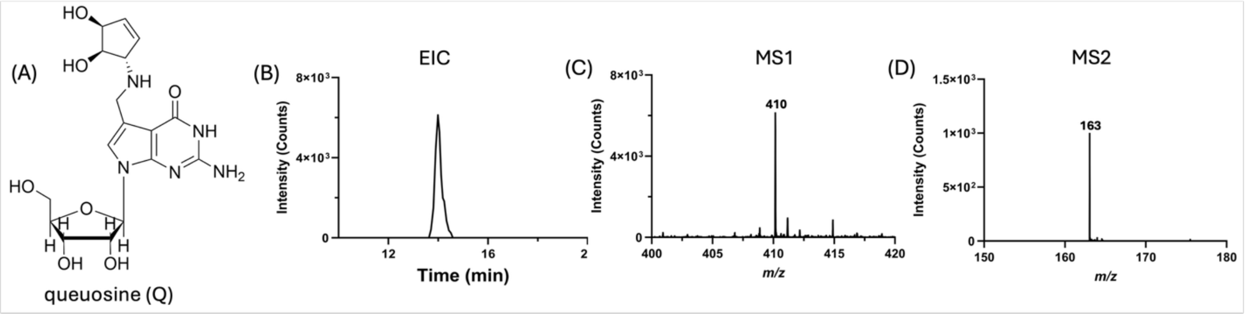


**Figure S1. Identification of queosine (Q) in total RNA extracts from neuronal tissue of *Aplysia* using LC-MS/MS.** (A) Chemical structure of Q. (B) Exemplary extracted ion chromatogram showing elution of Q. (C) MS spectrum of Q showing $[M+H]^+$ of 410.1675 m/z at 20 parts-per-million (ppm). (D) MS/MS spectrum of Q based on expected fragmentation patterns from collision-induced dissociation (CID). Expected MS and MS/MS ions were obtained from the MODOMICS database.

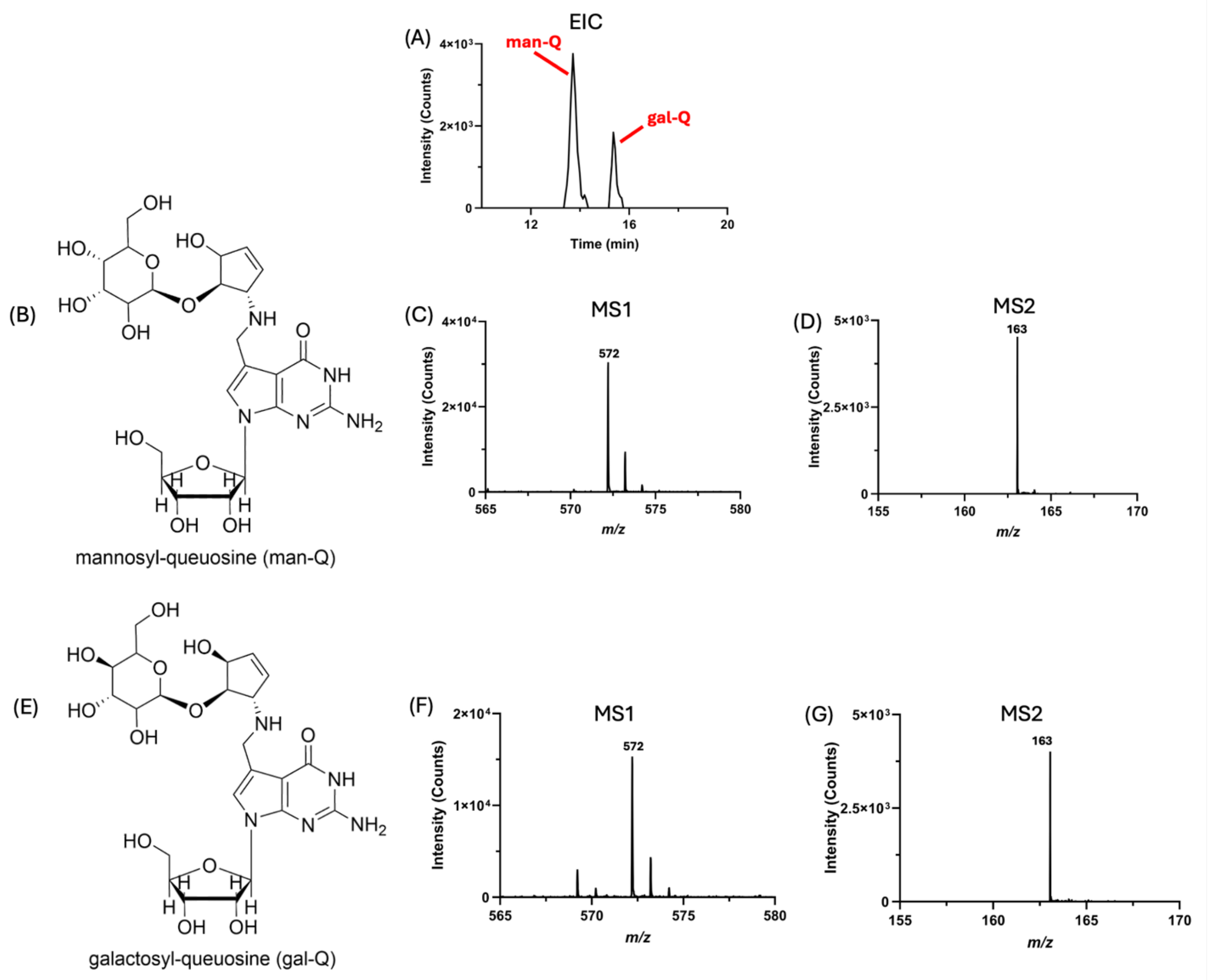


**Figure S2. Identification of mannosyl-queuosine (man-Q) and galactosyl-queuosine (gal-Q) in total RNA extracts from neuronal and non-neuronal tissue of *Aplysia* using LC-MS/MS.** (A) Extracted ion chromatogram showing elution of man-Q and gal-Q. Elution order was determined based off existing separation data using RP-HPLC.[95] (B) Chemical structure of man-Q. (C) MS and (D) MS/MS spectrum of man-Q showing $[M+H]^+$ of 572.2204 m/z at 20 ppm based on expected fragmentation patterns from collision-induced dissociation (CID). (E) Chemical structure of gal-Q. (F) MS and (G) MS/MS spectrum of gal-Q showing $[M+H]^+$ of 572.2204 m/z at 20 ppm with identical CID parameters. Expected MS and MS/MS ions were obtained from the MODOMICS database.

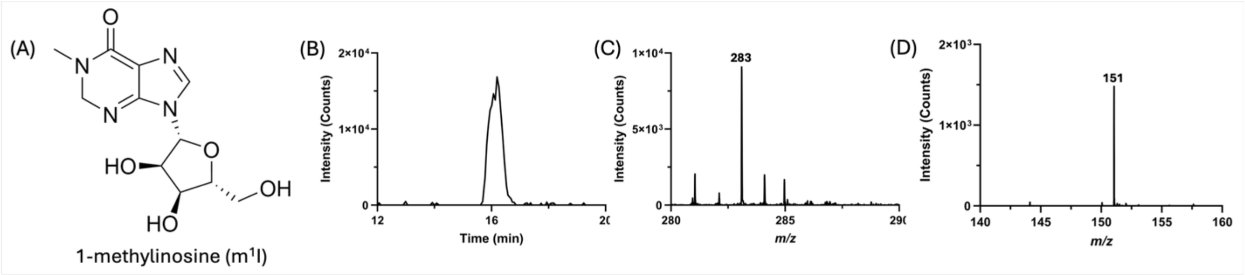


**Figure S3. Identification of 1-methylinosine ($m^1I$) in total RNA extracts from neuronal and non-neuronal tissue of *Aplysia* using LC-MS/MS. (A) Chemical structure of $m^1I$.** (B) Exemplary extracted ion chromatogram showing elution of $m^1I$. (C) MS spectrum of $m^1I$ showing $[M+H]^+$ of 283.1042 m/z at 20 ppm. (D) MS/MS spectrum of $m^1I$ based on expected fragmentation patterns from collision-induced dissociation (CID). Expected MS and MS/MS ions were obtained from the MODOMICS database.

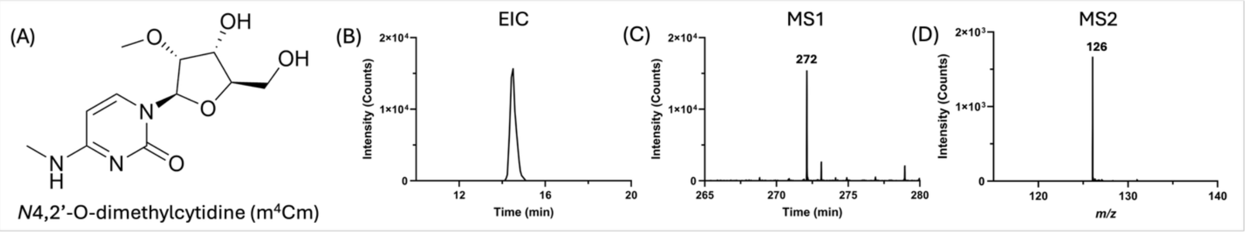


**Figure S4. Identification of *N*4,2-O-dimethylcytidine ($m^4Cm$) in total RNA extracts from neuronal and non-neuronal tissue of *Aplysia* using LC-MS/MS. (A) Chemical structure of $m^4Cm$.** (B) Exemplary extracted ion chromatogram showing elution of $m^4Cm$. (C) MS spectrum of $m^4Cm$ showing $[M+H]^+$ of 272.1246 m/z at 20 ppm. (D) MS/MS spectrum of $m^4Cm$ based on expected fragmentation patterns from collision-induced dissociation (CID). Expected MS and MS/MS ions were obtained from the MODOMICS database.

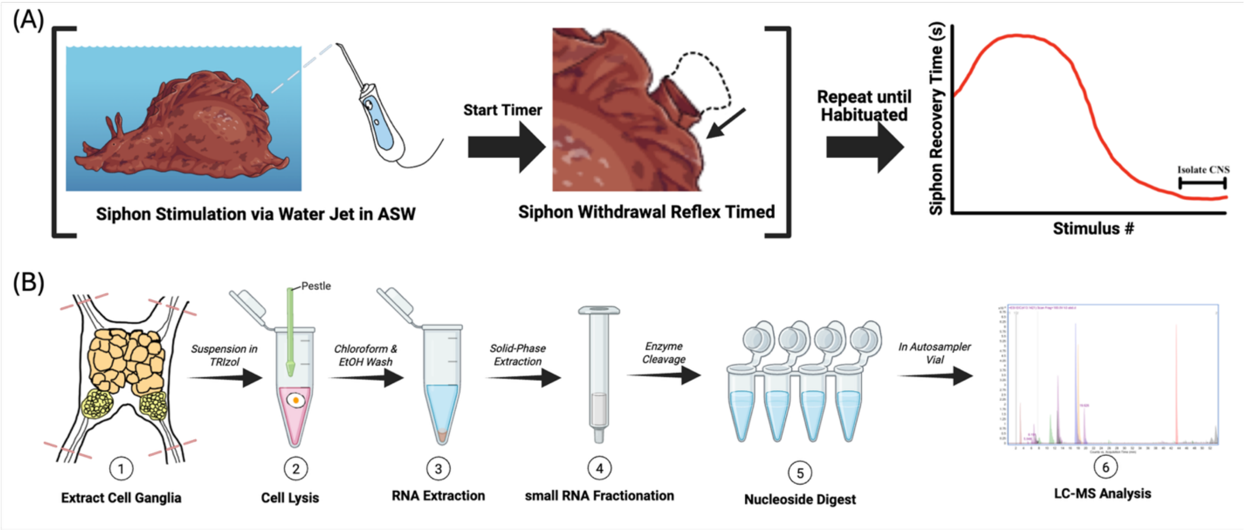


**Figure S5. Experimental workflow for behavioral training and RNA extraction in *Aplysia*.** (A) The animal's siphon-elicited siphon withdrawal reflex is timed (s) following consecutive exposure to mechanical stimuli via water jet. Successful habituation is indicated by visualizing significant depression of the reflex, which corresponds to a decrease in the siphon recovery time. (B) After successful habituation, total RNA from relevant ganglia is extracted, purified, and fractionated into small RNA (< 200 nt). Small RNA extracts are then enzymatically digested into nucleosides for LC-MS/MS analysis. Schematics were developed using BioRender.

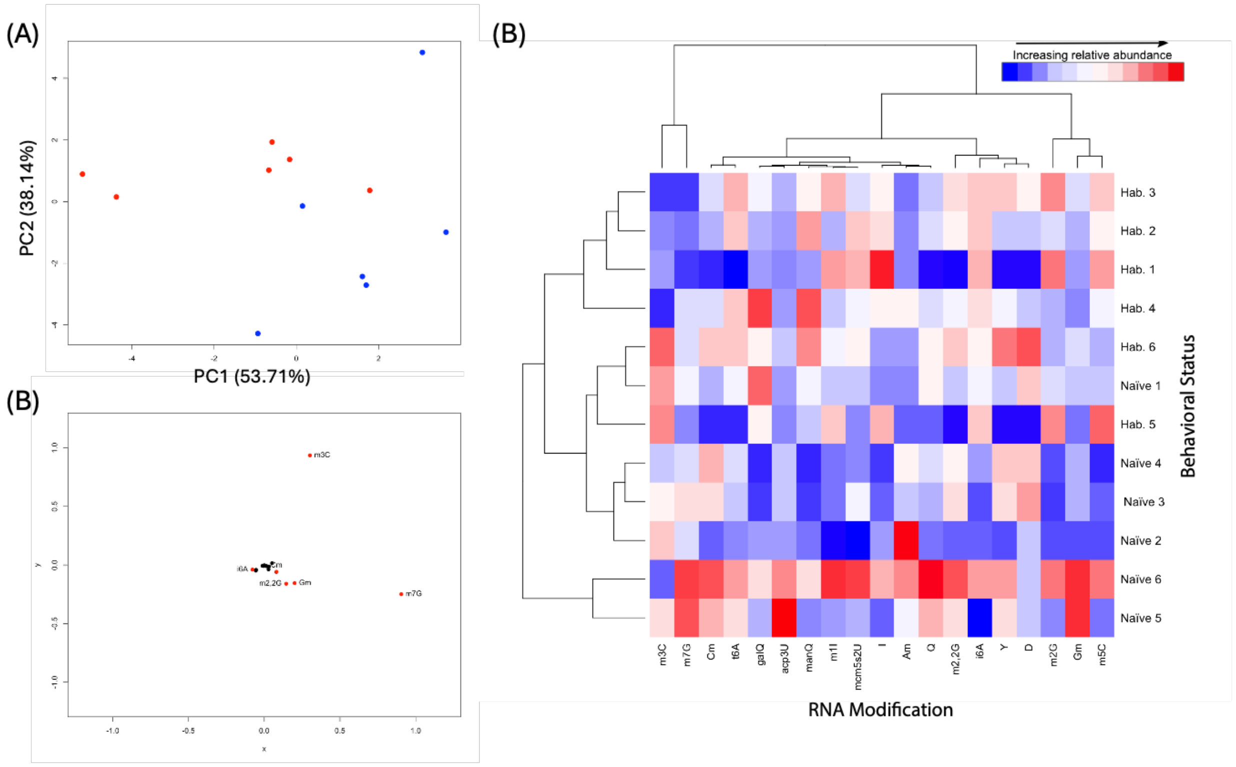


**Figure S6. Removal of m$^1$A from multivariate analysis improves unsupervised clustering between naive and habituated *Aplysia*.** (A) PCA of 19 neuronal modifications previously reported to be on tRNA detected in the abdominal ganglia of naïve and trained animals. (B) Loadings plot corresponding to the PCA in panel A. (C) Heat map diagram depicting hierarchical clustering of modified nucleosides based on normalized peak areas in abdominal ganglia of naïve and habituated animals without m$^1$A (n = 6).

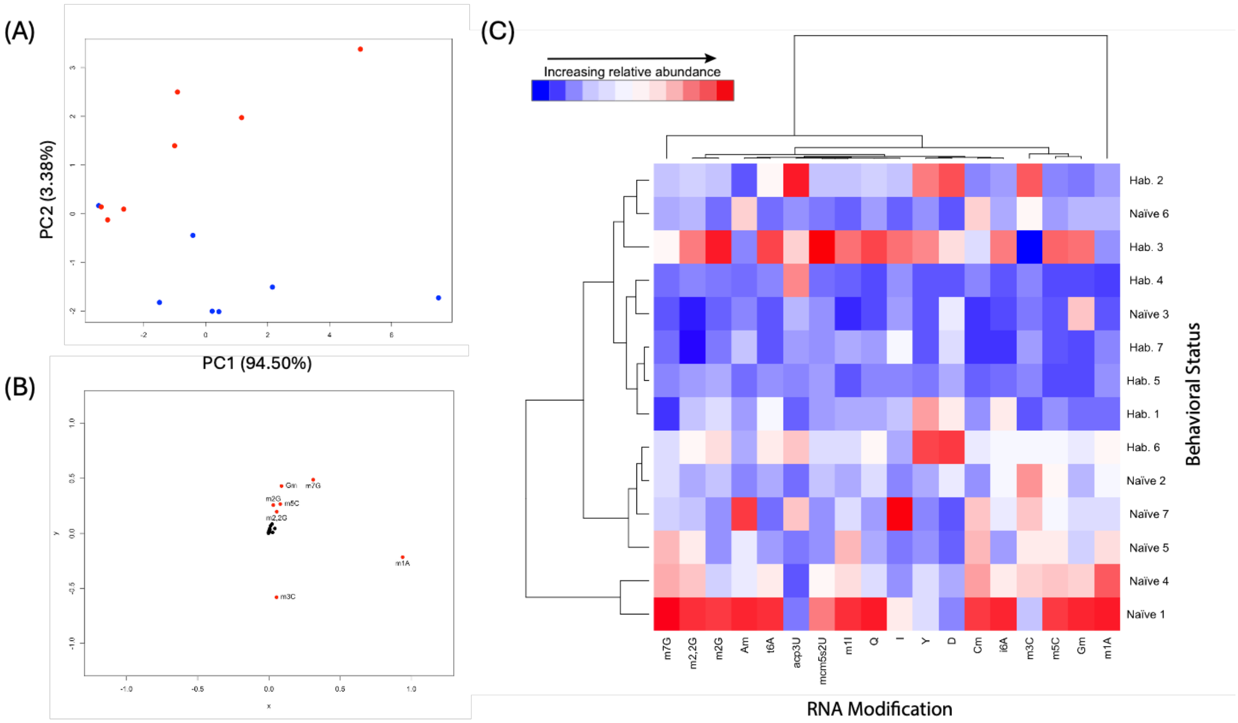


**Figure S7. Neuronal tRNA modifications in buccal ganglia subtly distinguish between naïve and habituated *Aplysia*.** (A) PCA of 18 neuronal tRNA modifications detected in buccal ganglia of naïve and trained animals. (B) Loadings plot corresponding to the PCA in panel A. (C) Heat map diagram depicting hierarchical clustering of modified nucleosides in buccal ganglia of naïve and habituated animals (n = 7).

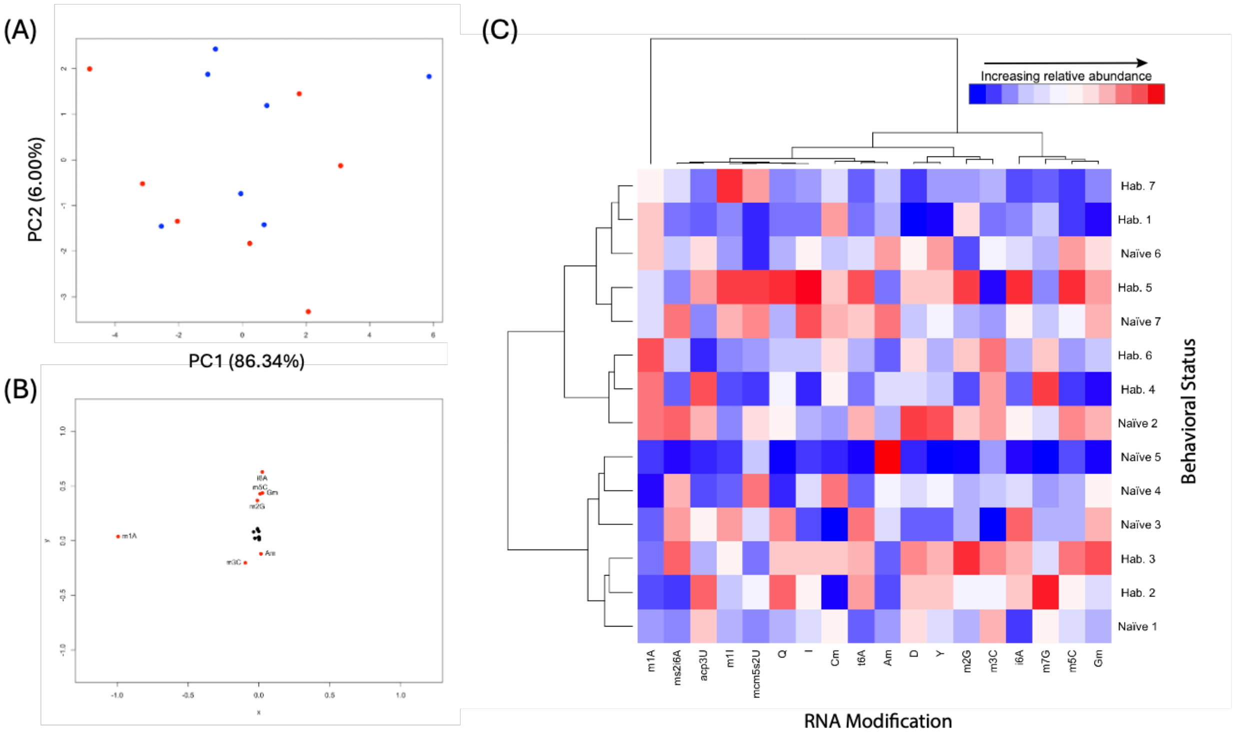


**Figure S8. Neuronal tRNA modifications in pedal ganglia do not distinguish between naïve and habituated *Aplysia*.** (A) PCA of 18 neuronal tRNA modifications detected in pedal ganglia of naïve and trained animals. (B) Loadings plot corresponding to the PCA in panel A. (C) Heat map diagram depicting hierarchical clustering of modified nucleosides in pedal ganglia of naïve and habituated animals (n = 7).

**Table S1. Successfully characterized modified ribonucleosides in 5 major ganglia and heart of *Aplysia californica* (n = 12).** Observed [M+H] and mass error show exemplary values from select replicates.

| Modification | Elution Order (RP-HPLC) | [M+H] observed | Mass error (ppm) | Abdominal | Pedal | Buccal | Heart |
|---|---|---|---|---|---|---|---|
| D | 1 | 247.0930 | -0.3 | 1,2,3 | 1,2,3 | 1,2,3 | 1,2,3 |
| Y | 2 | 245.0773 | 1.9 | 1,2,3 | 1,2,3 | 1,2,3 | 1,2,3 |
| $acp^3U$ | 3 | 346.125 | -5.5 | 1,2 | 1,2,3 | 1,2 | 1,2,3 |
| $m^3C$ | 4 | 258.109 | -3.8 | 1,2,3 | 1,2,3 | 1,2,3 | 1,2,3 |
| $m^1A$ | 5 | 282.1202 | 2.5 | 1,2,3 | 1,2,3 | 1,2,3 | 1,2,3 |
| $m^5C$ | 6 | 258.109 | -3 | 1,2,3 | 1,2,3 | 1,2,3 | 1,2,3 |
| Cm | 7 | 258.109 | -5.3 | 1,2,3 | 1,2,3 | 1,2,3 | 1,2,3 |
| $m^7G$ | 8 | 298.1151 | -2.7 | 1,2,3 | 1,2,3 | 1,2,3 | 1,2,3 |
| $m^4Cm$ | 9 | 272.1246 | -0.1 | 1,2,3 | 1,2,3 | 1,2,3 | 1,2,3 |
| I | 10 | 269.0886 | -4 | 1,2,3 | 1,2,3 | 1,2 | 1,2,3 |
| $m^1I$ | 11 | 283.1042 | 0.3 | 1,2,3 | 1,2 | 1,2 | 1,2,3 |
| Gm | 12 | 298.1151 | 7 | 1,2,3 | 1,2,3 | 1,2,3 | 1,2,3 |
| $m^2G$ | 13 | 298.1151 | 2.6 | 1,2,3 | 1,2,3 | 1,2,3 | 1,2,3 |
| Q | 14 | 410.1675 | -1.8 | 1,2,3 | 1,2,3 | 1,2 | 1,2,3 |
| manQ | 15 | 572.2204 | -3.3 | 1,2,3 | 1,2 | 1,2 | 1,2,3 |
| galQ | 16 | 572.2204 | -3.3 | 1,2,3 | 1,2,3 | 1,2 | 1,2,3 |
| $m^{2,2,7}G$ | 17 | 326.1464 | 0.9 | 1,2,3 | 1,2,3 | 1,2 | 1,2,3 |
| $m^{2,2}G$ | 18 | 312.1308 | 3.3 | 1,2,3 | 1,2,3 | 1,2,3 | 1,2,3 |
| Am | 19 | 282.1202 | 1.1 | 1,2,3 | 1,2,3 | 1,2,3 | 1,2,3 |
| $t^6A$ | 20 | 413.1421 | 1.1 | 1,2,3 | 1,2,3 | 1,2 | 1,2,3 |
| $m^6A$ | 21 | 282.1202 | 3.5 | 1,2,3 | 1,2,3 | 1,2,3 | 1,2,3 |
| $mcm^5s^2U$ | 22 | 333.0756 | -2.7 | 1,2,3 | 1,2 | 1,2 | 1,2,3 |
| $m^1Y/m^3Y$ | 23 | 259.093 | 5.5 | 1,2,3 | 1,2 | 1,2 | 1,2 |
| $m^{6,6}A$ | 24 | 296.1359 | 1.2 | 1,2,3 | 1,2,3 | 1,2,3 | 1,2,3 |
| $i^6A$ | 25 | 336.1672 | 4.6 | 1,2,3 | 1,2,3 | 1,2,3 | 1,2,3 |
| $ms^2i^6A$ | 26 | 382.1549 | -0.52 | 1,2,3 | 1,2 | 1,2 | 1,2,3 |

*Verified by: (1) elution order, (2) MS spectrum, (3) MS/MS spectrum*

**Table S2. Statistical analysis of relative peak areas of modified nucleosides detected by LC-QTOF-MS/MS from small RNA extracts of abdominal (n=6), buccal (n = 7), pedal (n = 7) ganglia and the heart (n = 7) of *Aplysia*.**

| Mod | i6A | m3C | D | Q | m2G | m5C | m1I | Cm | Am | I | m2,2G | mcm5s2U | acp3U | t6A | Gm | Y | m7G | m1A | | |
|---|---|---|---|---|---|---|---|---|---|---|---|---|---|---|---|---|---|---|---|---|
| **Abdominal 1** | 0.0173 | 0.1100 | 0.0176 | 0.0042 | 0.0347 | 0.0560 | 0.0021 | 0.0071 | 0.0027 | 0.0043 | 0.0208 | 0.0023 | 0.0001 | 0.0090 | 0.0522 | 0.0130 | 0.0943 | 0.7632 | | |
| **Abdominal 2** | 0.0171 | 0.0990 | 0.0142 | 0.0036 | 0.0304 | 0.0513 | 0.0017 | 0.0047 | 0.0057 | 0.0044 | 0.0140 | 0.0018 | 0.0007 | 0.0073 | 0.0446 | 0.0087 | 0.0883 | 0.7496 | | |
| **Abdominal 3** | 0.0163 | 0.0924 | 0.0187 | 0.0038 | 0.0297 | 0.0514 | 0.0019 | 0.0099 | 0.0032 | 0.0042 | 0.0247 | 0.0024 | 0.0001 | 0.0082 | 0.0506 | 0.0157 | 0.1013 | 0.7429 | | |
| **Abdominal 4** | 0.0168 | 0.0814 | 0.0173 | 0.0040 | 0.0298 | 0.0492 | 0.0019 | 0.0107 | 0.0037 | 0.0040 | 0.0237 | 0.0022 | 0.0010 | 0.0086 | 0.0507 | 0.0160 | 0.0914 | 0.7212 | | |
| **Abdominal 5** | 0.0184 | 0.0953 | 0.0144 | 0.0045 | 0.0319 | 0.0525 | 0.0020 | 0.0108 | 0.0035 | 0.0042 | 0.0257 | 0.0022 | 0.0034 | 0.0094 | 0.0703 | 0.0158 | 0.1326 | 0.8160 | | |
| **Abdominal 6** | 0.0222 | 0.0617 | 0.0147 | 0.0054 | 0.0404 | 0.0631 | 0.0027 | 0.0135 | 0.0045 | 0.0052 | 0.0333 | 0.0028 | 0.0033 | 0.0105 | 0.0705 | 0.0177 | 0.1396 | 0.8061 | | |
| **Buccal 1** | 0.0251 | 0.0836 | 0.0119 | 0.0073 | 0.0452 | 0.0810 | 0.0036 | 0.0270 | 0.0169 | 0.0054 | 0.0467 | 0.0033 | 0.0007 | 0.0136 | 0.1026 | 0.0114 | 0.2735 | 1.1914 | | |
| **Buccal 2** | 0.0178 | 0.1058 | 0.0145 | 0.0047 | 0.0290 | 0.0609 | 0.0022 | 0.0127 | 0.0039 | 0.0037 | 0.0259 | 0.0021 | 0.0005 | 0.0078 | 0.0656 | 0.0099 | 0.1793 | 0.9768 | | |
| **Buccal 3** | 0.0127 | 0.0739 | 0.0145 | 0.0031 | 0.0230 | 0.0453 | 0.0016 | 0.0078 | 0.0039 | 0.0039 | 0.0151 | 0.0017 | 0.0014 | 0.0063 | 0.0811 | 0.0071 | 0.1364 | 0.8302 | | |
| **Buccal 4** | 0.0187 | 0.0981 | 0.0130 | 0.0046 | 0.0297 | 0.0664 | 0.0027 | 0.0218 | 0.0074 | 0.0038 | 0.0351 | 0.0024 | 0.0002 | 0.0098 | 0.0790 | 0.0118 | 0.2153 | 1.1238 | | |
| **Buccal 5** | 0.0154 | 0.0933 | 0.0110 | 0.0042 | 0.0265 | 0.0622 | 0.0028 | 0.0205 | 0.0075 | 0.0035 | 0.0326 | 0.0017 | 0.0006 | 0.0077 | 0.0688 | 0.0093 | 0.2099 | 1.0109 | | |
| **Buccal 6** | 0.0156 | 0.0927 | 0.0105 | 0.0040 | 0.0242 | 0.0517 | 0.0018 | 0.0193 | 0.0096 | 0.0032 | 0.0265 | 0.0016 | 0.0009 | 0.0066 | 0.0646 | 0.0090 | 0.1616 | 0.9170 | | |
| **Buccal 7** | 0.0173 | 0.0997 | 0.0106 | 0.0040 | 0.0260 | 0.0604 | 0.0024 | 0.0200 | 0.0157 | 0.0087 | 0.0277 | 0.0017 | 0.0029 | 0.0068 | 0.0701 | 0.0089 | 0.1745 | 0.9532 | | |
| **Pedal 1** | 0.1056 | 0.0757 | 0.0414 | 0.0057 | 0.0559 | 0.1003 | 0.0037 | 0.0244 | 0.0164 | 0.0061 | 0.0621 | 0.0037 | 0.0110 | 0.0153 | 0.1052 | 0.0450 | 0.0971 | 0.4316 | | |
| **Pedal 2** | 0.1221 | 0.0773 | 0.0450 | 0.0060 | 0.0629 | 0.1084 | 0.0035 | 0.0195 | 0.0169 | 0.0058 | 0.0673 | 0.0041 | 0.0111 | 0.0177 | 0.1133 | 0.0493 | 0.0941 | 0.5364 | | |
| **Pedal 3** | 0.1321 | 0.0357 | 0.0379 | 0.0063 | 0.0563 | 0.0979 | 0.0045 | 0.0101 | 0.0182 | 0.0054 | 0.0725 | 0.0039 | 0.0107 | 0.0183 | 0.1137 | 0.0422 | 0.0918 | 0.4037 | | |
| **Pedal 4** | 0.1176 | 0.0608 | 0.0402 | 0.0053 | 0.0561 | 0.0976 | 0.0032 | 0.0306 | 0.0187 | 0.0061 | 0.0654 | 0.0044 | 0.0089 | 0.0152 | 0.1099 | 0.0455 | 0.0902 | 0.3669 | | |
| **Pedal 5** | 0.1014 | 0.0566 | 0.0365 | 0.0051 | 0.0445 | 0.0919 | 0.0030 | 0.0138 | 0.0281 | 0.0053 | 0.0468 | 0.0038 | 0.0084 | 0.0141 | 0.0935 | 0.0393 | 0.0776 | 0.3905 | | |
| **Pedal 6** | 0.1177 | 0.0650 | 0.0412 | 0.0057 | 0.0500 | 0.1071 | 0.0035 | 0.0211 | 0.0217 | 0.0062 | 0.0675 | 0.0034 | 0.0108 | 0.0166 | 0.1109 | 0.0476 | 0.0913 | 0.5007 | | |
| **Pedal 7** | 0.1216 | 0.0568 | 0.0400 | 0.0057 | 0.0563 | 0.1012 | 0.0044 | 0.0268 | 0.0232 | 0.0071 | 0.0694 | 0.0043 | 0.0095 | 0.0175 | 0.1133 | 0.0455 | 0.0928 | 0.4635 | | |
| **Heart 1** | 0.1691 | 0.0836 | 0.0737 | 0.0116 | 0.1568 | 0.1552 | 0.0053 | 0.0640 | 0.0133 | 0.0096 | 0.0785 | 0.0037 | 0.0225 | 0.0347 | 0.1434 | 0.0789 | 0.2090 | 0.7441 | | |
| **Heart 2** | 0.1699 | 0.0954 | 0.0788 | 0.0127 | 0.1198 | 0.1519 | 0.0062 | 0.0565 | 0.0093 | 0.0089 | 0.1311 | 0.0037 | 0.0245 | 0.0300 | 0.1321 | 0.0825 | 0.2166 | 0.7974 | | |
| **Heart 3** | 0.1525 | 0.1115 | 0.0761 | 0.0106 | 0.1313 | 0.1634 | 0.0056 | 0.0371 | 0.0097 | 0.0098 | 0.1086 | 0.0055 | 0.0275 | 0.0343 | 0.1453 | 0.0984 | 0.2269 | 0.6351 | | |
| **Heart 4** | 0.2079 | 0.0901 | 0.0754 | 0.0139 | 0.1714 | 0.1821 | 0.0070 | 0.0693 | 0.0105 | 0.0119 | 0.1474 | 0.0057 | 0.0275 | 0.0379 | 0.1662 | 0.0807 | 0.2306 | 0.9027 | | |
| **Heart 5** | 0.1740 | 0.0779 | 0.0735 | 0.0135 | 0.1535 | 0.1726 | 0.0066 | 0.0634 | 0.0149 | 0.0105 | 0.1257 | 0.0050 | 0.0245 | 0.0340 | 0.1562 | 0.0826 | 0.2182 | 0.7970 | | |
| **Heart 6** | 0.1670 | 0.0842 | 0.0682 | 0.0119 | 0.1407 | 0.1583 | 0.0056 | 0.0559 | 0.0107 | 0.0098 | 0.1181 | 0.0041 | 0.0224 | 0.0324 | 0.1418 | 0.0735 | 0.1841 | 0.7344 | | |
| **Heart 7** | 0.2030 | 0.1246 | 0.0951 | 0.0143 | 0.1712 | 0.1996 | 0.0073 | 0.0721 | 0.0122 | 0.0124 | 0.1508 | 0.0059 | 0.0315 | 0.0417 | 0.1767 | 0.0952 | 0.2328 | 0.8362 | | |
| **Mean (Abd)** | 0.0180 | 0.0900 | 0.0161 | 0.0042 | 0.0328 | 0.0539 | 0.0021 | 0.0095 | 0.0039 | 0.0044 | 0.0237 | 0.0023 | 0.0015 | 0.0088 | 0.0565 | 0.0145 | 0.1079 | 0.7665 | | |
| **Mean (Buc)** | 0.0175 | 0.0924 | 0.0123 | 0.0046 | 0.0291 | 0.0611 | 0.0024 | 0.0184 | 0.0093 | 0.0046 | 0.0299 | 0.0021 | 0.0010 | 0.0084 | 0.0760 | 0.0096 | 0.1929 | 1.0005 | | |
| **Mean (Ped)** | 0.1168 | 0.0611 | 0.0403 | 0.0057 | 0.0546 | 0.1006 | 0.0037 | 0.0209 | 0.0205 | 0.0060 | 0.0644 | 0.0039 | 0.0100 | 0.0164 | 0.1086 | 0.0449 | 0.0907 | 0.4419 | | |
| **Mean (Hrt)** | 0.1776 | 0.0953 | 0.0773 | 0.0126 | 0.1492 | 0.1690 | 0.0062 | 0.0598 | 0.0115 | 0.0104 | 0.1229 | 0.0048 | 0.0258 | 0.0350 | 0.1517 | 0.0845 | 0.2169 | 0.7781 | | |
| **SD (Abd)** | 0.0022 | 0.0167 | 0.0019 | 0.0006 | 0.0042 | 0.0050 | 0.0004 | 0.0031 | 0.0011 | 0.0004 | 0.0063 | 0.0004 | 0.0015 | 0.0011 | 0.0111 | 0.0032 | 0.0224 | 0.0372 | | |
| **SD (Buc)** | 0.0039 | 0.0107 | 0.0017 | 0.0013 | 0.0075 | 0.0113 | 0.0007 | 0.0063 | 0.0052 | 0.0020 | 0.0097 | 0.0006 | 0.0009 | 0.0026 | 0.0134 | 0.0016 | 0.0447 | 0.1228 | | |
| **SD (Ped)** | 0.0104 | 0.0140 | 0.0027 | 0.0004 | 0.0058 | 0.0057 | 0.0005 | 0.0072 | 0.0042 | 0.0006 | 0.0084 | 0.0004 | 0.0011 | 0.0015 | 0.0073 | 0.0033 | 0.0062 | 0.0615 | | |
| **SD (Hrt)** | 0.0202 | 0.0169 | 0.0085 | 0.0013 | 0.0196 | 0.0171 | 0.0008 | 0.0116 | 0.0020 | 0.0013 | 0.0247 | 0.0010 | 0.0033 | 0.0038 | 0.0155 | 0.0090 | 0.0167 | 0.0848 | **Average** | **SD** |
| **RSD (Abd)** | 12.018 | 18.522 | 11.980 | 15.077 | 12.669 | 9.3369 | 17.242 | 32.782 | 27.468 | 9.9898 | 26.711 | 15.5562 | 105.147 | 12.194 | 19.656 | 22.295 | 20.720 | 4.8555 | 21.9010 | 21.956 |
| **RSD (Buc)** | 22.116 | 11.565 | 14.212 | 28.841 | 25.745 | 18.440 | 27.513 | 34.223 | 56.347 | 42.484 | 32.458 | 29.1503 | 88.6863 | 31.009 | 17.585 | 16.769 | 23.160 | 12.276 | 29.5877 | 18.499 |
| **RSD (Ped)** | 8.9028 | 22.932 | 6.7857 | 7.0793 | 10.580 | 5.6913 | 14.689 | 34.354 | 20.379 | 10.232 | 13.062 | 9.0436 | 11.0330 | 9.3974 | 6.6887 | 7.3517 | 6.8148 | 13.914 | 12.1627 | 7.2762 |
| **RSD (Hrt)** | 11.370 | 17.711 | 11.018 | 10.542 | 13.127 | 10.113 | 12.563 | 19.484 | 17.693 | 12.457 | 20.084 | 19.9590 | 12.6274 | 10.868 | 10.241 | 10.598 | 7.7208 | 10.899 | 13.2819 | 3.8750 |

**Table S3:** Target mitochondrial tRNAs and their corresponding forward and reverse primer sequences.

| tRNA Isoacceptor | Forward Primer (5′ → 3′) | Reverse Primer (5′ → 3′) |
|---|---|---|
| tRNA-$Ser^{UCN}$ | AAT TTT TGT ACT ATA AGG TTT TGA A | GAA TTT TTC TCC CAT CAA GG |
| tRNA-$Tyr^{UAC}$ | TTT TGG TGG CTG ATT AAG G | AAT CCT AAG GGG TGA AAT CAC |
| tRNA-$Cys^{UGY}$ | TAA GTA GTA GTT TAG TAT AAA ACC | AAA AGT AGG AGC TAA GCT |
| tRNA-$Phe^{UUY}$ | TGA CAA ATA GCT TAA ATT TAA AGC ATA | TTA ACA AAA GGG CGG AGC |
| GAPDH | TCC ACT GGA GTC TTC ACA ACC | TGC AGG TCC TTG GTG TAC TTC |

**Table S4:** Relative Codon Usage of Target tRNAs in *Aplysia* Mitochondrial Genome

| mt-tRNA | Anticodon | Codon | Codon Usage Database (Frequency/10000) | CoCoPUTs (Frequency/1000) | Relative Ranking |
|---|---|---|---|---|---|
| Tyr | GUA | UAC | 9.4 | 9.36 | 3 |
| Cys | GCA | UGY | UGU: 12.2<br><u>UGC: 0.9</u> | UGU: 11.84<br><u>UGC: 1.10</u> | 4 |
| Phe | GAA | UUY | UUU: 63.7<br><u>UUC: 21.6</u> | UUU: 64.45<br><u>UUC: 21.48</u> | 1 |
| Ser | UGA | UCN | UCU: 39.5<br>UCC: 6.5<br><u>UCA: 11.9</u><br>UCG: 3.7 | UCU: 39.93<br>UCC: 6.61<br><u>UCA: 12.12</u><br>UCG: 3.86 | 2 |

*Underlined codons correspond to the specific anticodon that was observed for the isoacceptor.*
